# High burden and seasonal variation of paediatric scabies and pyoderma prevalence in The Gambia: a cross-sectional study

**DOI:** 10.1101/591537

**Authors:** Edwin P. Armitage, Elina Senghore, Saffiatou Darboe, Momodou Barry, Janko Camara, Sulayman Bah, Michael Marks, Carla Cerami, Anna Roca, Martin Antonio, Claire E. Turner, Thushan I. de Silva

**Affiliations:** Medical Research Council Unit The Gambia at The London School of Hygiene & Tropical Medicine, PO Box 273, Banjul, The Gambia; Department of Clinical Research, Faculty of Infectious and Tropical Diseases, London School of Hygiene & Tropical Medicine, London, UK; Department of Molecular Biology & Biotechnology, The Florey Institute, University of Sheffield, Sheffield, UK; Department of Infection, Immunity and Cardiovascular Diseases, The Florey Institute, University of Sheffield, Sheffield, UK

**Keywords:** scabies, pyoderma, group A streptococcus / *Streptococcus pyogenes*, rheumatic heart disease, The Gambia

## Abstract

**Background:** Scabies is a WHO neglected tropical disease common in children in low-and middle-income countries. Excoriation of scabies lesions can lead to secondary pyoderma infection, most commonly by *Staphyloccocus aureus* and *Streptococcus pyogenes* (group A streptococcus, GAS), with the latter linked to acute post-streptococcal glomerulonephritis (APSGN) and potentially rheumatic heart disease (RHD). There is a paucity of data on the prevalence of these skin infections and their bacterial aetiology from Africa.

**Materials/methods:** A cross-sectional study, conducted over a four-month period that included the dry and rainy season, was conducted to determine the prevalence of common skin infections in Sukuta, a peri-urban settlement in western Gambia, in children <5 years. Swabs from pyoderma lesions were cultured for *S. aureus* and GAS.

**Results:** Of 1441 children examined, 15.9% had scabies (95% CI 12.2-20.4), 17.4% had pyoderma (95% CI 10.4-27.7) and 9.7% had fungal infections (95% CI 6.6-14.0). Scabies were significantly associated with pyoderma (aOR 2.74, 95% CI 1.61-4.67). Of 250 pyoderma swabs, 80.8% were culture-positive for *S. aureus*, and 50.8% for GAS. Participants examined after the first rains were significantly more likely to have pyoderma than those examined before (aRR 2.42, 95% CI 1.38-4.23), whereas no difference in scabies prevalence was seen (aRR 1.08, 95% CI 0.70-1.67). Swab positivity was not affected by the season.

**Conclusions:** High prevalence of scabies and pyoderma were observed. Pyoderma increased significantly during rainy season. Given the high prevalence of GAS pyoderma among children, further research on the association with RHD in West Africa is warranted.

**Summary:** This cross-sectional study of skin infections in The Gambia revealed prevalence of scabies, pyoderma and fungal infections in children <5 years of 15.9%, 17.4% and 9.7% respectively, with increased bacterial skin infections in the rainy season.

## Background

*Streptococcus pyogenes* (group A streptococcus, GAS) is a pathogen that causes a wide spectrum of disease, from superficial skin infections through to invasive sepsis and streptococcal toxic shock syndrome [1]. It continues to cause a significant burden of morbidity and mortality globally, particularly in low-and middle-income countries (LMIC) [2]. It is also associated with autoimmune-mediated, post-infective sequelae including acute post-streptococcal glomerulonephritis (APSGN) and acute rheumatic fever (ARF) leading to chronic rheumatic heart disease (RHD) [3]. RHD has largely disappeared from high-income countries except in neglected groups such as Aboriginal Australians [3, 4], but remains a significant problem in LMIC including those in sub-Saharan Africa (sSA) [5]. The classical understanding of ARF involves an acute GAS pharyngitis infection with specific “rheumatogenic” GAS *emm* serotypes [6, 7]. It is, however, increasingly thought that this explanation is incomplete, with a diversity of *emm* types and GAS skin infections potentially playing an important role in LMIC [8, 9]. This is supported by data from countries where GAS pyoderma and RHD are both highly prevalent, but pharyngitis and GAS pharyngeal carriage are low [3, 10, 11]. A significant proportion of GAS skin infections are attributable to secondary bacterial infection of scabies lesions, a WHO neglected tropical disease, known to be particularly prevalent in young children living in poverty-related conditions in tropical countries [12-14]. Despite this, there are few data sources on scabies epidemiology for sSA [15-20].

Pyoderma (defined as any infection of the skin involving pus) is common in LMIC independently of its association with scabies [12], with over 162 million children estimated to be suffering from impetigo globally [21]. Epidemiological and microbiological data on pyoderma in Africa is scarce [3, 22-24]. Due to importance of GAS as a pathogen in sSA and M-protein type-specific vaccines in development, a registry of GAS infections in Africa has been initiated to address the paucity of microbiological data [25]. Superficial fungal infections are also common globally, particularly in children in LMIC. The burden in Africa is heterogeneous, with prevalence in schoolchildren ranging from 10-80% [26].

In The Gambia, the prevalence of these common skin infections and the bacterial aetiology of pyoderma in children are unknown. To provide preliminary data, we performed a cross-sectional study to determine prevalence of scabies, pyoderma and fungal skin infections in children aged <5 years in a peri-urban community in western Gambia. We also aimed to characterise the microbiological aetiology of pyoderma in this setting.

## Methods

### Setting

The Gambia is the smallest country by area in mainland Africa, with a population of 1.9 million [27]. It was ranked 174^th^ in the United Nations Human Development Index in 2017 [28]. The annual seasonal climate consists of a long dry season from November to May and a short rainy season between June and October. Sukuta is an area within the West Coast Region peri-urban conurbation, with a population of 47,048 in 2013, including 7,234 children aged under 5 years [27].

### Study design and sampling

A cross-sectional, population-based study was conducted in Sukuta, in children aged <5 years, to determine the prevalence of scabies, pyoderma and fungal infections. Prior to the study, geospatial census data from 2013 was used to divide Sukuta into 37 geographical clusters of approximately 100 households (Figure S1). A one-stage, random cluster sampling method was used, sampling clusters in a randomly generated order. All households in each selected cluster were approached for participation. Eligible participants were children <5 years of age sleeping in that household. Participants were examined over a four-month period between May and September 2018.

### Diagnosis and training

Study nurses underwent a 2-day training led by a physician, to introduce basic dermatology skills, recognise common paediatric skin complaints, and instruct on use of an adapted IMCI algorithm (Figure S2) [29]. Scabies cases were diagnosed clinically based on the presence of pruritus and papules in a typical distribution. Pyoderma was defined as any skin lesion with evidence of pus or crusts. Infected scabies was diagnosed when scabies was present with evidence of inflammation and pyoderma in the same distribution. Fungal (dermatophyte) skin infection was diagnosed on the basis of the presence of round or oval flat scaly patches, with features typical of tinea infection. Where pyoderma was diagnosed, a wound swab was taken. All skin conditions identified were managed appropriately according to the algorithm and treatment guidelines (Figure S2 and Table S1).

### Algorithm validity

To determine the sensitivity and specificity of the algorithm used, a subset of participants were selected opportunistically and re-examined by a general physician with experience of diagnosing paediatric dermatological conditions in The Gambia, blinded to the nurse’s diagnosis. A sample size of 123 was calculated to be sufficient to detect a sensitivity of 80% compared to the gold standard examination with a precision of 10% assuming a prevalence of 50% of each condition.

### Swab collection and bacteriological methods

Wound swabs were collected using Nylon^®^ flocked swabs stored in liquid Amies transport medium and transported in a cold box to the MRC Unit The Gambia at LSHTM (MRCG) for same-day culturing on 5% sheep’s blood agar and MacConkey agar incubated overnight. Purity plates were done for mixed infection. Identification by catalase and confirmation for either *S*. *aureus* or GAS was done using the Remel™ Staphaurex™ Plus and Streptex™ latex agglutination tests, respectively. Antibiotic sensitivity pattern was obtained using disc diffusion according to CLSI methods [30].

### Investigation of seasonality

Data collection spanned both the dry and rainy season (defined as after the 26^th^ of June 2018 when the first rains of the year occurred), allowing for comparison between the prevalence, and the proportion of pyoderma caused by *S. aureus* and GAS between these two periods. In addition, the first cluster sampled was subsequently resampled at the end of the study in order to directly compare prevalence in the same population before and after the start of the rainy season.

To further investigate seasonal change in pyoderma in The Gambia, we also leveraged the presence of existing medical records from a rural primary healthcare clinic run by MRCG to perform a post-hoc analysis. The MRC Keneba clinic, in West Kiang, provides free primary care and has been collecting electronic medical records since 2009 [31]. Records on the number of presentations per month of <5 year olds for skin complaints (including impetigo, cutaneous abscess, furuncle or carbuncle, and cellulitis) between 2011 and 2018 were interrogated to ascertain if seasonal variation was observed.

### Data collection and statistical analysis

A questionnaire was delivered to participant’s parents to collect information on socio-demographics and risk factors for skin diseases. Data were collected on tablet computers using REDCap™ electronic data capture tools hosted at MRCG [32]. Data were analysed using Stata^®^ version 15.1. The cluster random sampling method was corrected for using the *svyset* command. Adjusted binomial exact confidence intervals were calculated for prevalence estimates. Associations between socio-demographic and other risk factors and skin infections were investigated using multivariable logistic regression models. Variables and category levels were selected for inclusion using forwards and backwards stepwise regression, without *svyset* correction, using a likelihood ratio test significance level of p<0.2. The multivariable models were then rerun with *svyset* correction to obtain adjusted odds ratios (aOR). All socio-demographics and risk factors were categorical variables, except household size, which was continuous. The population attributable risk (PAR) [33] for scabies as a cause of pyoderma was calculated. The adjusted prevalence ratio (aPR) for the prevalence of skin conditions before and after the start of the rainy season were estimated using a multivariable Poisson regression model, adjusting for socio-demographics, except when comparing the first cluster before and after the rains, which was unadjusted. Sensitivity, specificity and kappa statistic were calculated to determine the accuracy of the diagnostic algorithm compared to the diagnosis reached by the physician. Significance was set at p<0.05.

### Ethical considerations

Ethical approval for the study was provided by The Gambia Government/MRC Joint Ethics Committee (SCC1587). Written or thumb-printed informed consent was obtained from a parent for all participants included in the prevalence study prior to involvement.

## Results

### Prevalence of skin infections

A total of 1441 participants from 9 clusters were examined between May and September 2018. The mean number of participants per cluster was 160.1 (range 62-306). Overall, 47.9% of participants were male and the mean age was 28.9 (SD 16.7) months. Scabies prevalence was 15.9% (95% CI 12.2-20.4), pyoderma prevalence was 17.4% (95% CI 10.4-27.7), and fungal infection prevalence was 9.7% (95% CI 6.6-14.0). The prevalence of scabies, pyoderma and fungal infections by socio-demographic characteristics is presented in Table 1.

**Table 1.**
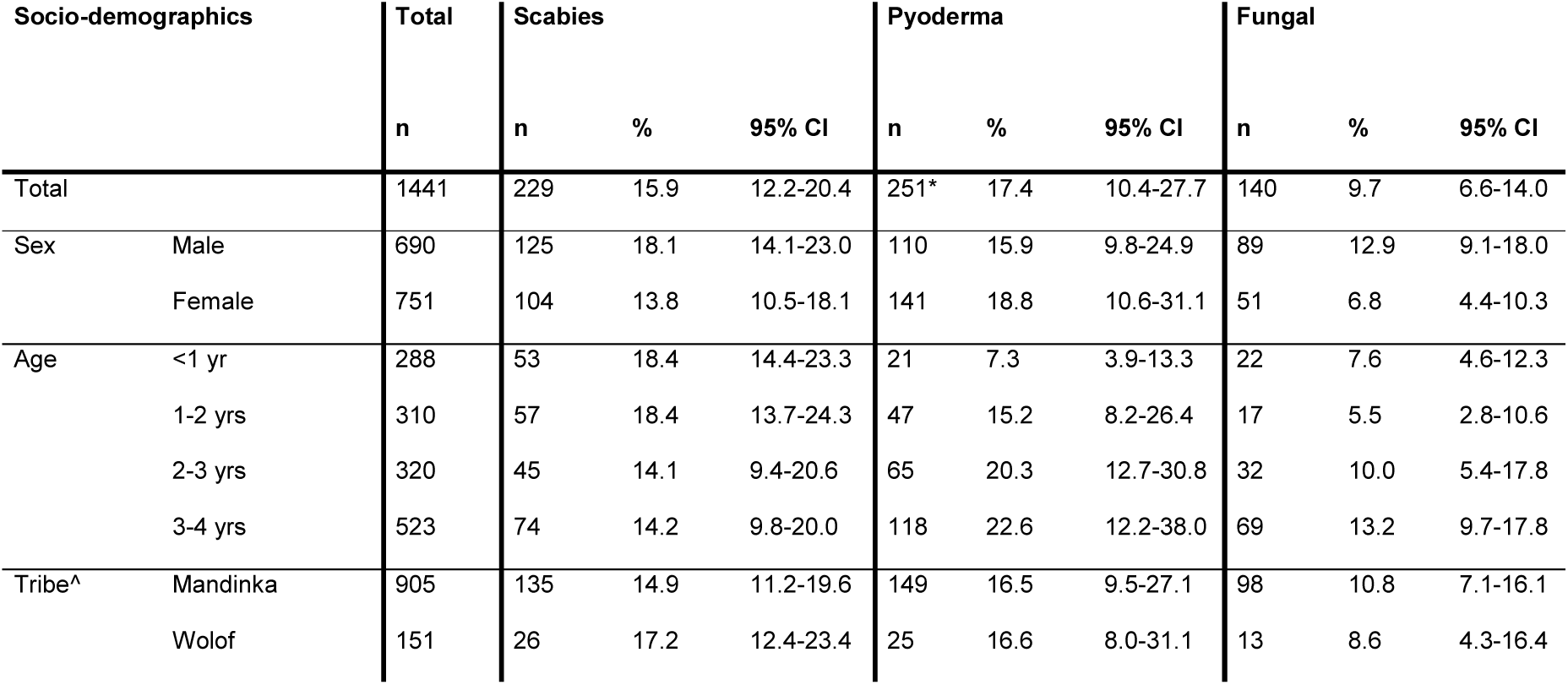
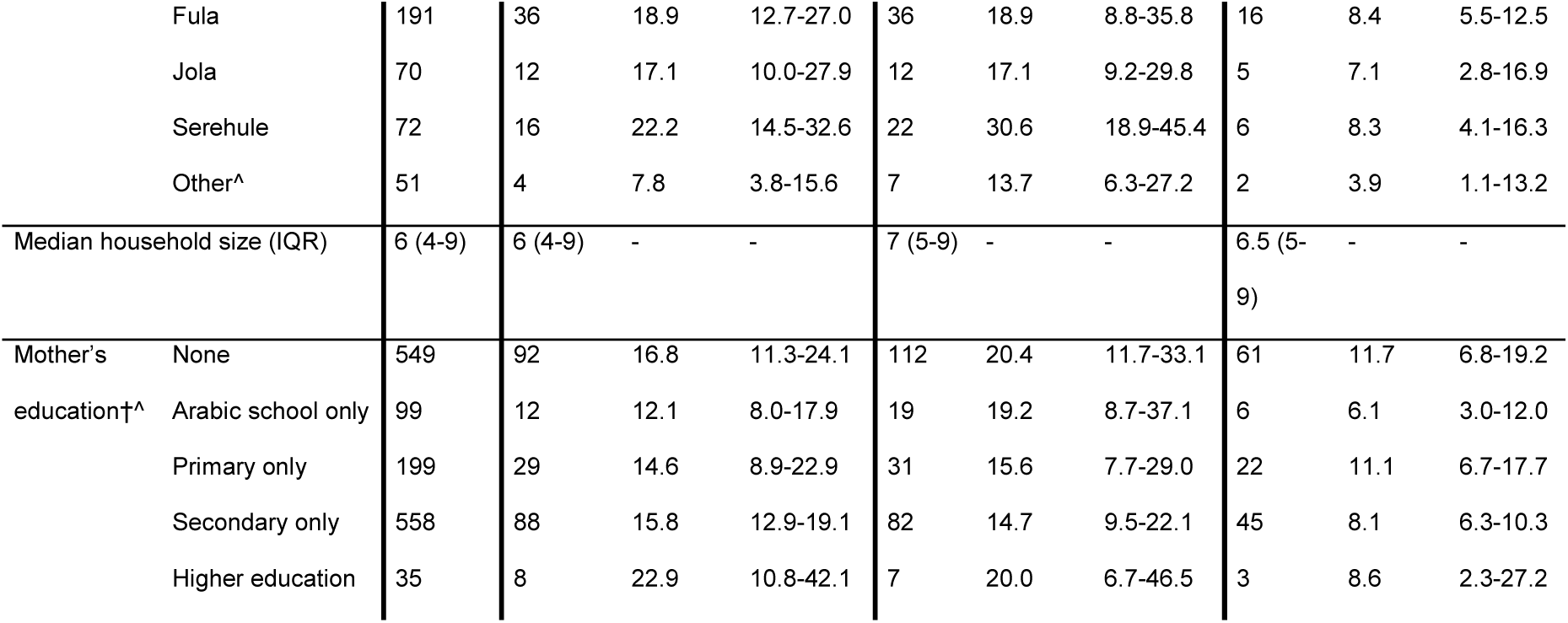
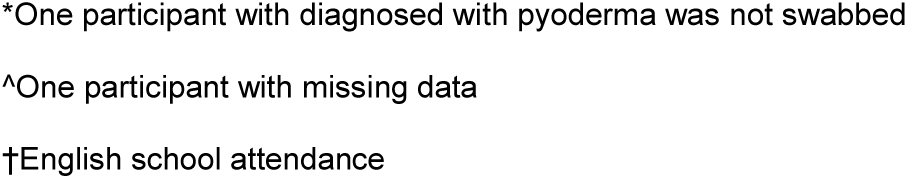
Prevalence of skin infections by socio-demographics. Confidence intervals are corrected for cluster sampling design. *One participant with diagnosed with pyoderma was not swabbed ^One participant with missing data †English school attendance

### Associations with socio-demographic and other risk factors

Table 2 shows the aOR for skin infection presence by socio-demographic and other risk factors. The odds of scabies and fungal infections were significantly lower in females than males (aOR 0.70, 95% CI 0.61-0.82 and aOR 0.44, 95% CI 0.32-0.61 respectively). Pyoderma increased with age (aOR 3.13 of pyoderma in 3-4 years compared to <1 year, 95% CI 2.00-4.88) and was higher in Serehule children compared to Mandinka children (aOR 1.99, 95% CI 1.16-3.41), and increased with household size (aOR 1.03, 95% CI 1.01-1.06). Various behavioural risk factors were identified, including lower odds of scabies in those whose clothes are always ironed (aOR 0.26, 95% CI 0.07-0.98), and higher odds of fungal infections in those not wearing freshly washed clothes every day (aOR 13.5, 95% CI 3.22-26.60). Current breastfeeding was protective against fungal infections, but increased the odds of scabies (aOR 1.67, 95% CI 1.12-2.49). A history of previous skin infections increased the odds of all three infections. Results from the forwards and backwards stepwise regression procedure are presented in Tables S2-S5.

**Table 2.** Adjusted odds ratios for socio-demographic and other risk factors potentially associated with skin infections. Reported values are from multivariable logistic regression models, including only variables and category levels selected by stepwise regression with an elimination level of p>0.2. For all three skin infections, forwards and backwards stepwise regression produced the same final model (Tables S4 and S5). Sex and age group were included in all models, and correction for cluster sampling design was done on the final models. Data from univariable logistic regression models are shown in Table S2. ref = reference category used; aOR = adjusted odds ratio; *significant at p<0.05; **significant at p<0.001; †English school attendance

|  |  | Scabies |  |  | Pyoderma |  |  | Fungal |  |  |
| --- | --- | --- | --- | --- | --- | --- | --- | --- | --- | --- |
|  |  | aOR | p value | 95% CIs | aOR | p value | 95% CIs | aOR | p value | 95% CIs |
| Sex | Male | ref |  |  | ref |  |  | ref |  |  |
|  | Female | 0.70 | 0.001* | 0.61-0.82 | 1.22 | 0.201 | 0.88-1.70 | 0.44 | <0.001** | 0.32-0.61 |
| Age category | <1 year | ref |  |  | ref |  |  | ref |  |  |
|  | 1-2 years | 0.91 | 0.634 | 0.60-1.40 | 1.98 | 0.086 | 0.89-4.43 | 0.48 | 0.006* | 0.30-0.76 |
|  | 2-3 years | 0.81 | 0.529 | 0.38-1.72 | 2.85 | <0.001** | 1.90-4.26 | 0.55 | 0.026* | 0.33-0.91 |
|  | 3-4 years | 0.87 | 0.670 | 0.42-1.79 | 3.13 | <0.001** | 2.00-4.88 | 0.79 | 0.509 | 0.36-1.74 |
| Tribe | Mandinka | ref |  |  | ref |  |  | ref |  |  |
|  | Serehule | 1.72 | 0.044* | 1.02-2.90 | 1.99 | 0.018* | 1.16-3.41 |  |  |  |
|  | Other | 0.42 | 0.020* | 0.21-0.84 |  |  |  | 0.39 | 0.105 | 0.12-1.28 |
| Mean household size |  |  |  |  | 1.03 | 0.014* | 1.01-1.06 |  |  |  |
| Mother's education† | None | ref |  |  |  |  |  | ref |  |  |
|  | Arabic school only | 0.60 | 0.007* | 0.43-0.83 |  |  |  | 0.48 | 0.040* | 0.24-0.96 |
|  | Secondary only |  |  |  |  |  |  | 0.69 | 0.126 | 0.42-1.14 |
| Currently breastfeeding | No | ref |  |  |  |  |  | ref |  |  |
|  | Yes | 1.67 | 0.017* | 1.12-2.49 |  |  |  | 0.52 | 0.043* | 0.27-0.98 |
| Low birth weight (<2.5kg) | No |  |  |  |  |  |  | ref |  |  |
|  | Yes |  |  |  |  |  |  | 1.80 | 0.006* | 1.25-2.59 |
| Water source | Tap |  |  |  | ref |  |  |  |  |  |
|  | Well |  |  |  | 1.41 | 0.434 | 0.54-3.73 |  |  |  |
| Clean clothes | Every day |  |  |  |  |  |  | ref |  |  |
|  | Not every day |  |  |  |  |  |  | 13.50 | 0.003* | 3.22-56.60 |
| Clothes ironed | Never | ref |  |  |  |  |  |  |  |  |
|  | Always | 0.26 | 0.048* | 0.07-0.98 |  |  |  |  |  |  |
| Handwashing | No | ref |  |  |  |  |  |  |  |  |
| area in compound | Yes | 0.74 | 0.107 | 0.50-1.09 |  |  |  |  |  |  |
| Open fire in | No | ref |  |  | ref |  |  |  |  |  |
| compound | Yes | 1.51 | 0.021* | 1.09-2.10 | 1.40 | 0.018* | 1.08-1.82 |  |  |  |
| Previous skin | None | ref |  |  | ref |  |  | ref |  |  |
| infection | One | 3.09 | <0.001** | 2.19-4.64 | 1.94 | 0.002* | 1.40-2.69 | 2.34 | 0.005* | 1.41-3.87 |
|  | More than one | 4.51 | <0.001** | 2.63-7.73 | 2.34 | 0.002* | 1.53-3.57 | 2.14 | 0.072 | 0.92-5.02 |
| History of burn | No | ref |  |  |  |  |  |  |  |  |
|  | Yes | 0.65 | 0.154 | 0.35-1.22 |  |  |  |  |  |  |
| History of | No |  |  |  | ref |  |  |  |  |  |
| malnutrition | Yes |  |  |  | 0.67 | 0.243 | 0.32-1.40 |  |  |  |
| History of | No | ref |  |  |  |  |  |  |  |  |
| nutritional | Yes | 0.32 | 0.170 | 0.06-1.83 |  |  |  |  |  |  |
| supplementation |  |  |  |  |  |  |  |  |  |  |

The presence of pyoderma was significantly associated with scabies infestation (aOR 2.74, 95% CI 1.61-4.67) as shown in Table 3. The PAR of scabies as a cause of pyoderma was 14.7% (95% CI 5.1-23.3).

**Table 3.** Adjusted odds ratios and population attributable risk percentage for scabies as a cause of pyoderma, *S. aureus* positive pyoderma and GAS positive pyoderma. Reported values are from a multivariable logistic regression model adjusting for sex, age group, tribe, household size and mother’s education, following correction for cluster sampling design. ref = reference category used; aOR = adjusted odds ratio; *significant at p<0.05; PAR = population attributable risk

|  |  | Scabies n (%) |  | aOR (p value, 95% CI) | PAR % (95% CI) |
| --- | --- | --- | --- | --- | --- |
|  |  | No | Yes |  |  |
| All pyoderma | No | 1031 (85.1) | 159 (69.4) | ref |  |
|  | Yes | 181 (14.9) | 70 (30.6) | 2.74* (0.002, 1.61-4.67) | 14.7 (5.1-23.3) |
| *S. aureus* positive pyoderma | No | 34 (18.9) | 14 (20.0) | ref |  |
|  | Yes | 146 (81.1) | 56 (80.0) | 1.08 (0.775, 0.61-1.91) | 0.4 (-2.7-3.4) |
| GAS positive pyoderma | No | 88 (48.9) | 35 (50.0) | ref |  |
|  | Yes | 92 (51.1) | 35 (50.0) | 1.26 (0.179, 0.88-1.81) | 2.8 (-1.7-7.1) |

### Pyoderma swab culture results

Pyoderma swabs were taken from 250 participants (one participant diagnosed with pyoderma was not swabbed). Overall, either *S. aureus* or GAS was cultured from 90.0% of swabs, with 80.8% positive for *S. aureus*, and 50.8% for GAS (41.6% were positive for both). Additionally, two samples were positive for other beta-haemolytic streptococci (one group D *Streptococcus* and one non-groupable *Streptococcus* species). The proportion of GAS and *S. aureus* causing pyoderma were similar regardless of scabies infection (Table 3). All GAS isolates were sensitive to penicillin, while 99% of *S. aureus* isolates were methicillin-sensitive. Antibiotic sensitivities of isolates are presented in Figures S3 and S4.

### Seasonal variation of skin infection prevalence

The respective prevalence of scabies, pyoderma and fungal infection were 15.3%, 8.9% and 14.4% before the start of the rainy season, compared to 16.3%, 23.1% and 6.6% after. The prevalence of scabies did not change significantly (aPR 1.08, 95% CI 0.70-1.67), whereas pyoderma prevalence significantly increased following the start of the rains (aPR 2.42, 95% CI 1.39-4.23). Fungal infection prevalence fell significantly (aPR 0.44, 95% CI 0.32-0.60) (Table S6). The PAR of scabies as a cause of pyoderma was 18.5% (95% CI 4.1-30.8) before the start of the rainy season compared to 13.6% (95% CI −0.7-25.8) afterwards. The prevalence of scabies, pyoderma and fungal skin infections by week sampled are presented in Figure 1. The increase in pyoderma prevalence during the rainy season was confirmed by resampling of the first cluster sampled (7.9% vs. 21.7%, PR 2.74, 95% CI 1.23-6.12). The start of the rains did not significantly affect the proportion of *S. aureus* or GAS detected (Table S6).

**Figure 1.**
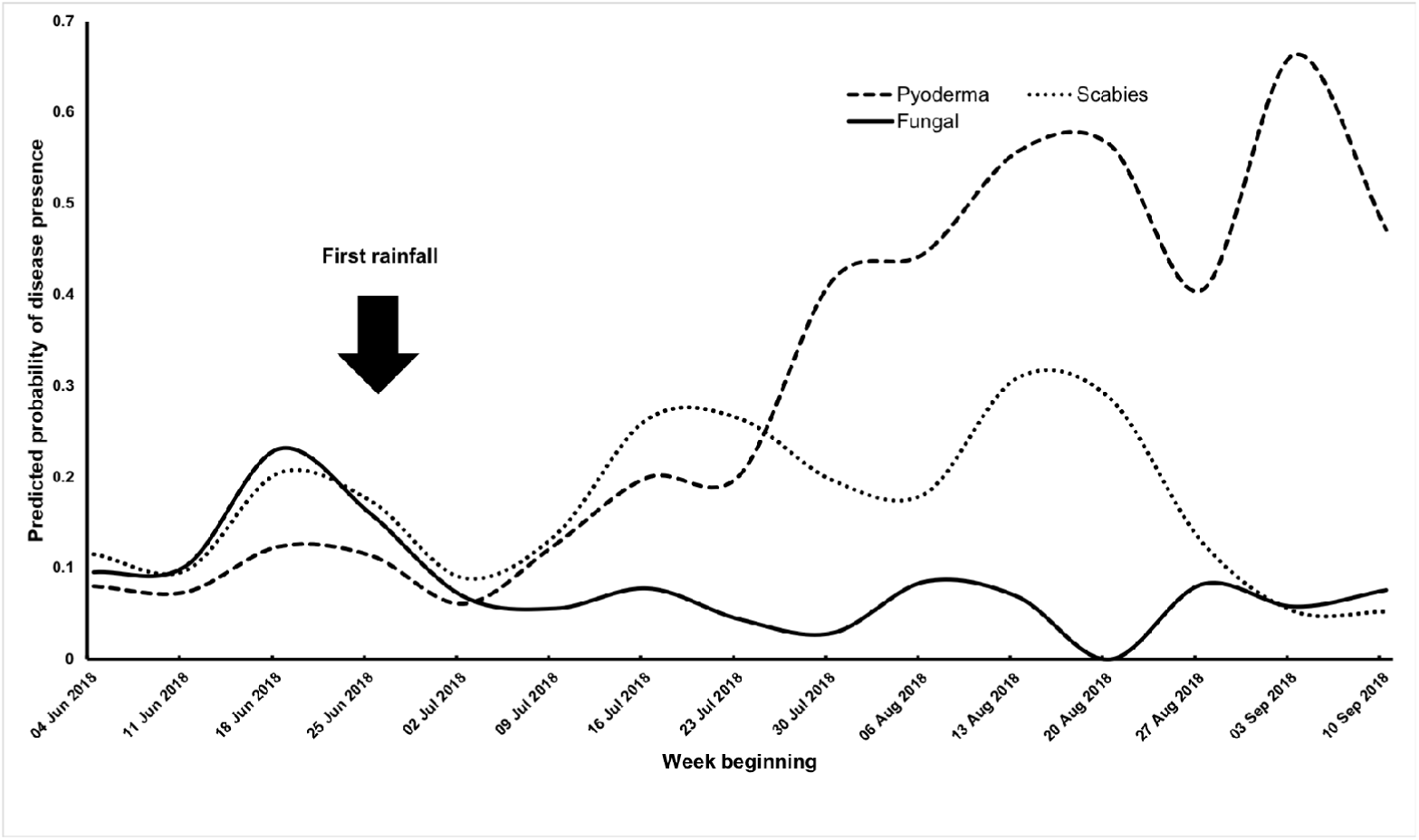
Predicted probabilities of skin infection prevalence by week examined.

**Figure 2.**
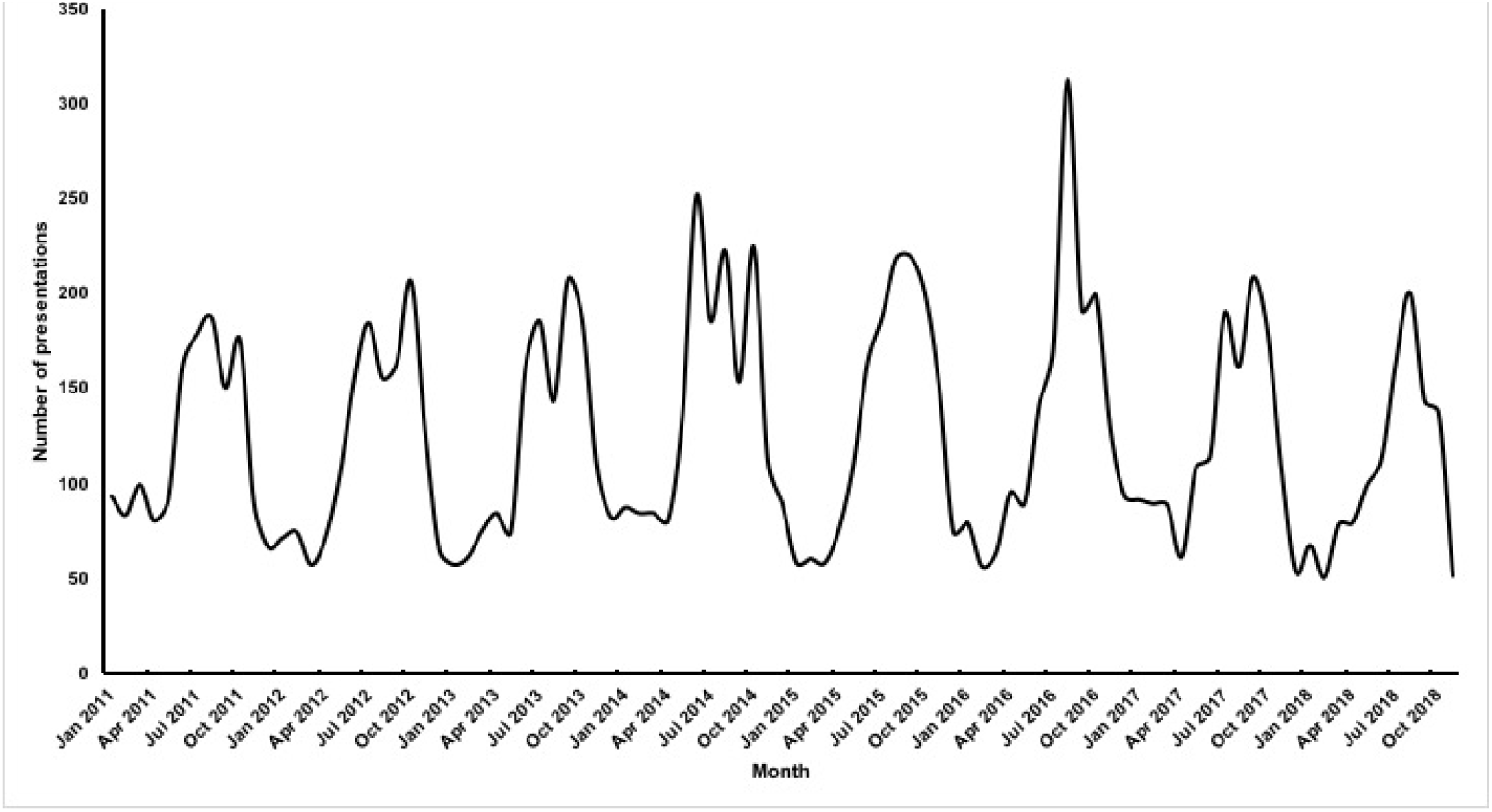
Number of skin complaint presentations in under 5s at MRCG Keneba clinic by month 2011-2018.

The data obtained from the MRCG Keneba clinic records of presentations per month of children <5 for all skin complaints are presented in Figure S3. The number of presentations peaked each year between June and December, coinciding with the rainy season.

### Sensitivity and specificity of the diagnostic algorithm

Diagnosis by nursing staff using the algorithm (Figure S2) was 97.1% sensitive and 96.6% specific for the diagnosis of pyoderma, and 83.3% sensitive and 97.0% specific for non-infected scabies. Sensitivity and specificity for the pyoderma and scabies co-infection were 81.3% and 97.2% respectively. Sensitivity of detection of fungal infections was lower at 66.7%, but specificity was 95.8%. Full results including the kappa statistic for inter-rater agreement are presented in Table S7.

## Discussion

We found a high burden of skin infections among Gambian children and that pyoderma increased significantly during the rainy season. The scabies prevalence we observed of 15.9% is consistent with other studies from sSA [15, 16, 18-20], but the overall prevalence of pyoderma of 17.4% is higher than the estimated median impetigo prevalence of 7% for Africa [21]. Studies in other settings have found a strong association between scabies and pyoderma, with one study in Fiji finding a PAR of scabies as a cause of impetigo of 93.1% [10]. We observed a PAR of 14.7%, indicating that scabies may play a less significant role in pyoderma rates in this setting. Interventions to reduce scabies, shown to be effective elsewhere [34-36], may still be worthwhile given the relatively high scabies prevalence.

The 8.9% pyoderma prevalence in children examined before the start of the rainy season was consistent with previous estimates for Africa [21]. However, in those examined during the rainy season, the prevalence rate was significantly higher (23.1%). This effect was confirmed in the cluster that was sampled both before and after the start of the rainy season. Furthermore, a strong seasonal trend was seen at the rural MRCG Keneba clinic in paediatric skin presentations between 2011 and 2018, which supports the hypothesis that bacterial skin infections are seasonal in The Gambia, a finding also observed in 1980 [17].

Very little microbiology data exist from pyoderma cases in Africa [3, 21]. By taking swabs from pyoderma lesions, we were able to provide robust estimates for the prevalence of *S. aureus* and GAS pyoderma. 50.8% of swabs were positive for GAS, which is lower than the global median of 74% noted previously from the limited heterogeneous studies available [21]. However, with the high pyoderma prevalence during the rainy season, this still indicates a significant exposure of children <5 to GAS and *S. aureus* annually. Better understanding the carriage and transmission of these organisms within households and communities in The Gambia are required to design preventative interventions and reduce the disease burden.

The reason for the seasonal increase in skin infections is not clear, but may reflect behavioural changes between seasons or environmental changes enhancing bacterial proliferation on skin. Indoor space can be limited within households, and during the rainy season people may spend more time in overcrowded indoor areas, increasing skin-to-skin contact. This could partly explain the observed increase, although scabies prevalence did not increase correspondingly as would be expected. Further studies are required to explore the exact mechanism for this observation. We also observed associations between increased age, household size, history of previous infections, and certain tribal groups with higher pyoderma rates, some of which have been observed elsewhere [13, 37]. These suggest that individual’s behaviour may also be important in pyoderma transmission.

Additionally, associations between scabies and fungal infections and other risk factors were seen. Current breastfeeding increased the odds of scabies, which may reflect higher risk due to greater skin contact with mothers who may be the source of infection, while ironing of clothes reduced the odds of scabies, which could reduce transmission of mites through clothes. The largest effect-size was seen in participants not wearing freshly washed clothes every day, which increased the odds of fungal infections. These may represent potential targets for behavioural interventions to reduce these skin infections.

Nurses were used for the diagnosis of the skin conditions rather than clinicians as nurses are commonly the primary healthcare providers in LMIC. The sensitivity and specificity of the algorithm used were consistent with the validation study [29], confirming that such algorithms for nurses can be effective diagnostic tools.

Our study has several limitations. We sampled from one peri-urban population, so our findings may not be generalisable, particularly to rural settings, where the majority of Gambians live. No up-to-date sampling frame was available, we therefore used a random cluster sampling method which reduced the precision of the prevalence confidence intervals. The study was conducted over four months, so we were unable to determine whether the observed seasonal variation is cyclical on an annual basis or changes year-to-year. We were only able to resample one cluster to compare prevalence directly, and given that prevalence varied by cluster, we should interpret the results cautiously. Additionally, the associations observed between risk factors and skin infections should be interpreted with caution as question responses were open to recall bias, and there may have been residual confounding factors not measured. Finally, the subset used for the algorithm validation sub-study were selected opportunistically, which may have been open to selection bias. Despite these limitations, this study provides good baseline data for the prevalence of these common skin infections in a country where no other recent data exists and also provides valuable microbiological data on skin pathogens, which is lacking globally [1-3].

## Conclusions

Skin infections may be overlooked, particularly in LMIC where there are other pressing health concerns, but with such high prevalence, especially of pyoderma during the rainy season, they may represent a crucial early exposure to potentially serious pathogens. The high incidence of invasive *S. aureus* infections in Gambian children is increasingly recognised and can be triggered by skin infections [38]. The exact mechanism by which RHD follow GAS infection is still not understood, but repeated GAS skin infections may be important [39]. It is plausible that exposure to GAS through the skin at a young age may be a step in the aetiology of RHD [8, 9], particularly as GAS pyoderma is known to trigger the better understood, immune-complex mediated APSGN, which is also epidemiologically linked to scabies [23, 40]. Increased research GAS carriage, transmission dynamics, and correlates of protection from disease is therefore warranted in African settings. Treating skin infections for their own sake is justification enough, but tackling these as an upstream target may also be a cost-effective way to make a significant impact on more serious conditions.

## Funding

This work was supported by a HEFCE/ODA grant from The University of Sheffield (155123). TdS in funded by a Wellcome Trust Intermediate Clinical Fellowship award [110058/ Z/15/Z]. CET is a Royal Society & Wellcome Trust Sir Henry Dale Fellow [208765/Z/17/Z]. MM is supported by an NIHR Clinical Lectureship. Research at the MRC Unit The Gambia at LSHTM is jointly funded by the UK Medical Research Council (MRC) and the UK Department for International Development (DFID) under the MRC/DFID Concordat agreement and is also part of the EDCTP2 programme supported by the European Union.

## Conflicts of interest

None.

## Acknowledgements

We acknowledge the hard work of the field workers and nurses and the MRCG clinical microbiology laboratory team. We are grateful to the study participants and their families and the wider Sukuta community for their involvement and assistance. We would like to thank The MRCG Keneba Clinic, Abdoulie Faal and Andrew Prentice for access to the KEMRES clinical data.

