## Appendix for "High burden and seasonal variation of paediatric scabies and pyoderma prevalence in The Gambia: a cross-sectional study"

**Supplementary Appendix**

**Figure S1. The Sukuta area divided into 37 geographical clusters with approximately equal number of compounds from 2013 geospatial census data, used for one-stage cluster sampling method.**

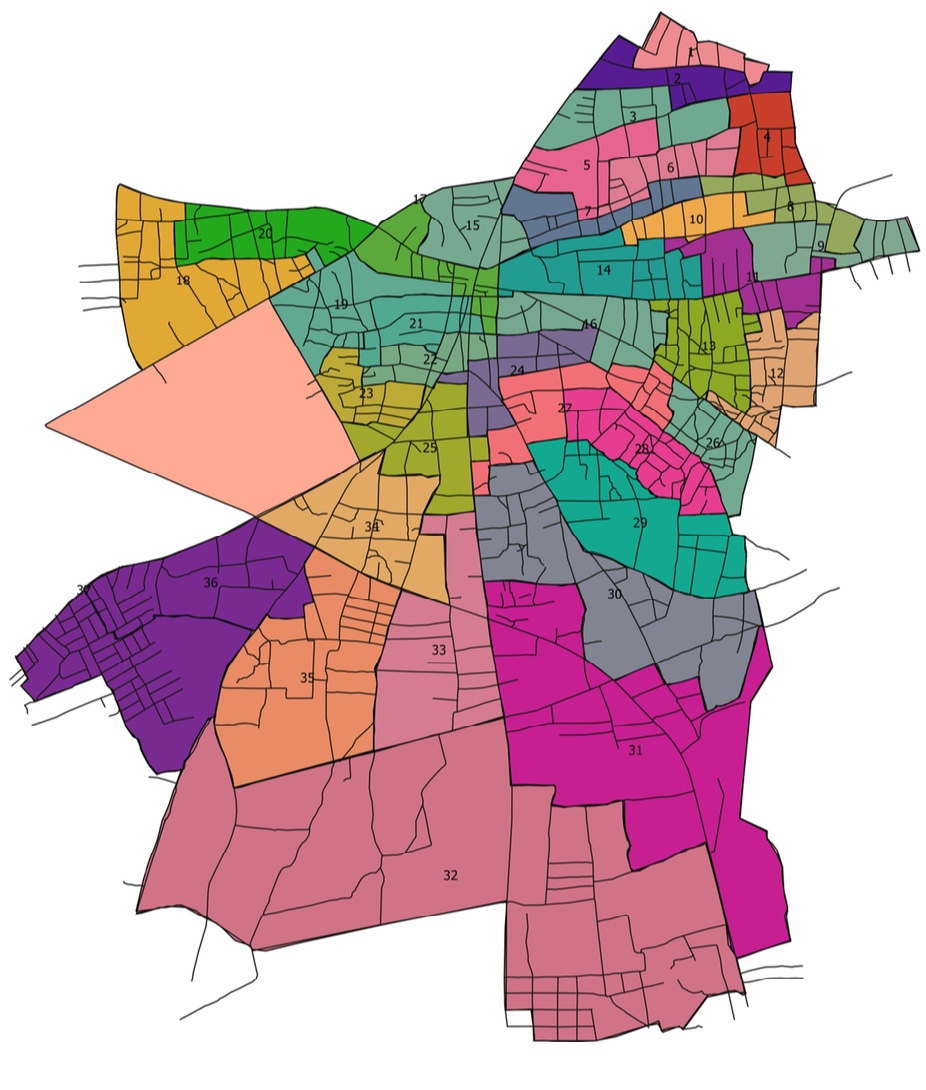

**Figure S2.** **Diagnostic algorithm for common skin conditions adapted from IMCI algorithm designed and validated by Steer *et al.* (2009)**
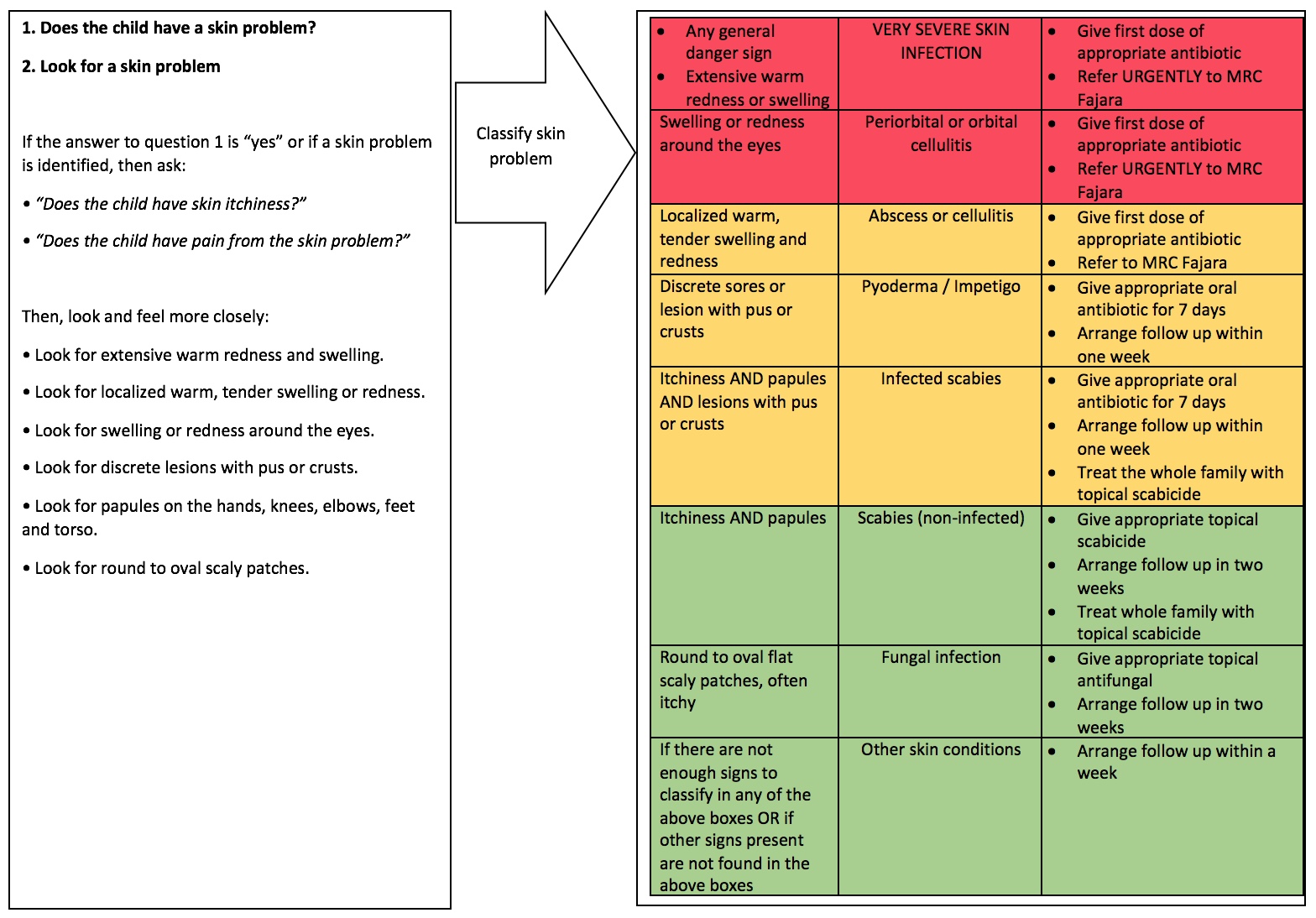

**Table S1. Treatment guidelines used for common skin conditions**. *Where participants were diagnosed with infected scabies they were not treated with benzyl benzoate until after the bacterial infection had resolved **Where participant were diagnosed with non-infected or infected scabies all close family contacts were also treated.

| **Diagnosis** | **Drug** | **Age** | **Dosage and course length** |
| --- | --- | --- | --- |
| Very severe skin infection / Periorbital or orbital cellulitis / Abscess or cellulitis | Cloxacillin syrup | Under 28 days | 25mg/kg stat oral dose before referral |
|  |  | 1 month to 2 years | 62.5mg stat oral dose  before referral |
|  |  | 2 - 4 years | 125mg stat oral dose  before referral |
| *Alternative (if penicillin allergy)* | Azithromycin syrup | All | 12mg/kg once a day for 5 days |
| Pyoderma / Impetigo / Infected scabies* | Cloxacillin syrup | Under 7 days | 25mg/kg twice a day for one week |
|  |  | 7-20 days | 25mg/kg three times a day for one week |
|  |  | 21-28 days | 25mg/kg four times a day for one week |
|  |  | 1 month to 2 years | 62.5mg four times a day for one week |
|  |  | 2 - 4 years | 125mg four times a day for one week |
| *Alternative (if penicillin allergy)* | Azithromycin syrup | All | 12mg/kg once a day for 5 days |
| Scabies (non-infected)** | Benzyl benzoate 25% w/v emulsion | Under 1 year | Dilute with 3 parts water, apply once and wait for 24 hours before washing |
|  |  | 1 - 18 years | Dilute with equal parts water, apply once and wait for 24 hours before washing |
|  |  | Adults (family members) | Do not dilute, apply once and wait for hour hours before washing |
| Fungal infection | Clotrimazole 1% cream | All | Apply twice a day for two weeks |

**Table S2**. **Odds ratios for socio-demographic and other risk factors potentially associated with skin infections in univariable logistic regression models**. All values were corrected for cluster sampling design.

ref = reference category used; NA = regression analysis not possible due to too few participants; *significant at p<0.05; **significant at p<0.001

|  |  | **Scabies** | | | **Pyoderma** | | | **Fungal** | | |
| --- | --- | --- | --- | --- | --- | --- | --- | --- | --- | --- |
|  |  | **OR** | **p value** | **95% CIs** | **OR** | **p value** | **95% CIs** | **OR** | **p value** | **95% CIs** |
| Sex | Male | ref |  |  | ref |  |  | ref |  |  |
|  | Female | 0.73 | <0.001** | 0.65-0.81 | 1.22 | 0.226 | 0.86-1.73 | 0.49 | <0.001** | 0.39-0.63 |
| Age category | <1 year | ref |  |  | ref |  |  | ref |  |  |
|  | 1-2 years | 1.00 | 0.995 | 0.68-1.46 | 2.27 | 0.043* | 1.03-5.00 | 0.70 | 0.192 | 0.40-1.24 |
|  | 2-3 years | 0.73 | 0.151 | 0.46-1.16 | 3.24 | <0.001** | 2.21-4.75 | 1.34 | 0.144 | 0.88-2.04 |
|  | 3-4 years | 0.73 | 0.169 | 0.45-1.18 | 3.70 | <0.001** | 2.39-5.73 | 1.84 | 0.005* | 1.28-2.64 |
| Tribe | Mandinka | ref |  |  | ref |  |  | ref |  |  |
|  | Wolof | 1.19 | 0.252 | 0.86-1.63 | 1.01 | 0.971 | 0.67-1.52 | 0.78 | 0.348 | 0.43-1.40 |
|  | Fula | 1.32 | 0.277 | 0.76-2.31 | 1.18 | 0.644 | 0.54-2.59 | 0.75 | 0.128 | 0.51-1.11 |
|  | Jola | 1.18 | 0.469 | 0.71-1.95 | 1.05 | 0.901 | 0.44-2.52 | 0.63 | 0.256 | 0.27-1.50 |
|  | Serehule | 1.63 | 0.069 | 0.95-2.79 | 2.23 | 0.015* | 1.22-4.08 | 0.75 | 0.458 | 0.32-1.76 |
|  | Other | 0.49 | 0.084 | 0.21-1.13 | 0.81 | 0.624 | 0.31-2.13 | 0.34 | 0.036 | 0.12-0.91 |
| Mean household size | | 1.02 | 0.209 | 0.99-1.04 | 1.03 | 0.029* | 1.00-1.05 | 1.02 | 0.036* | 1.00-1.04 |
| Mother’s education† | None | ref |  |  | ref |  |  | ref |  |  |
|  | Arabic school only | 0.69 | 0.048* | 0.47-1.00 | 0.93 | 0.829 | 0.42-2.04 | 0.49 | 0.074 | 0.22-1.09 |
|  | Primary only | 0.85 | 0.577 | 0.44-1.63 | 0.72 | 0.280 | 0.37-1.38 | 0.94 | 0.795 | 0.53-1.57 |
|  | Secondary only | 0.93 | 0.675 | 0.63-1.37 | 0.67 | 0.061 | 0.44-1.02 | 0.66 | 0.061 | 0.43-1.03 |
|  | Higher education | 1.47 | 0.197 | 0.78-2.77 | 0.98 | 0.942 | 0.46-2.09 | 0.71 | 0.610 | 0.16-3.14 |
| Currently breastfeeding | No | ref |  |  | ref |  |  | ref |  |  |
|  | Yes | 1.47 | 0.033* | 1.04-2.08 | 0.42 | <0.001** | 0.32-0.54 | 0.49 | 0.003* | 0.33-0.72 |
| Low birth weight (<2.5kg) | No | ref |  |  | ref |  |  | ref |  |  |
|  | Yes | 1.17 | 0.227 | 0.88-1.56 | 1.09 | 0.707 | 0.66-1.79 | 1.33 | 0.128 | 0.90-1.98 |
|  | Unknown | 1.08 | 0.562 | 0.80-1.45 | 1.50 | 0.002* | 1.22-1.85 | 1.19 | 0.309 | 0.83-1.70 |
| Water source | Tap | ref |  |  | ref |  |  | ref |  |  |
|  | Borehole | 1.30 | 0.390 | 0.67-2.52 | 0.99 | 0.988 | 0.28-3.49 | 1.00 | 0.988 | 0.55-1.80 |
|  | Well | 1.36 | 0.350 | 0.67-2.76 | 1.48 | 0.371 | 0.57-3.87 | 1.11 | 0.645 | 0.66-1.87 |
| Water distance | Inside compound | ref |  |  | ref |  |  | ref |  |  |
|  | <5 mins away | 0.51 | 0.122 | 0.20-1.25 | 1.04 | 0.910 | 0.50-2.17 | 1.21 | 0.499 | 0.65-2.23 |
|  | 5-10 mins away | 1.05 | 0.823 | 0.63-1.77 | 1.27 | 0.199 | 0.86-1.89 | 0.86 | 0.039 | 0.74-0.99 |
|  | >10 mins away | 1.20 | 0.714 | 0.40-3.62 | 0.65 | 0.379 | 0.23-1.88 | 0.66 | 0.296 | 0.278-1.56 |
| Full body wash | Every day | ref |  |  | ref |  |  | ref |  |  |
|  | Not every day | 0.75 | 0.686 | 0.16-3.55 | 0.68 | 0.605 | 0.13-3.62 | NA | NA | NA |
| Clean clothes | Every day | ref |  |  | ref |  |  | ref |  |  |
|  | Not every day | 0.48 | 0.374 | 0.08-2.91 | NA | NA | NA | 13.63 | 0.003* | 3.23-57.48 |
| Clothes ironed | Never | ref |  |  | ref |  |  | ref |  |  |
|  | Sometimes | 0.88 | 0.176 | 0.73-1.07 | 1.40 | 0.005* | 1.14-1.83 | 1.59 | 0.039* | 1.03-2.44 |
|  | Always | 0.18 | 0.011* | 0.06-0.60 | NA | NA | NA | 2.00 | 0.180 | 0.67-5.96 |
| Handwashing area in compound | No | ref |  |  | ref |  |  | ref |  |  |
|  | Yes | 0.71 | 0.046* | 0.50-0.99 | 0.78 | 0.106 | 0.57-1.07 | 0.87 | 0.352 | 0.64-1.20 |
| Open fire in compound | No | ref |  |  | ref |  |  | ref |  |  |
|  | Yes | 1.49 | 0.004* | 1.18-1.88 | 1.35 | 0.034* | 1.03-1.77 | 1.34 | 0.347 | 0.68-2.64 |
| Previous skin infection | None | ref |  |  | ref |  |  | ref |  |  |
|  | One | 2.68 | <0.001** | 1.99-3.60 | 2.21 | 0.001* | 1.58-3.10 | 2.32 | 0.002* | 1.49-3.63 |
|  | More than one | 3.33 | 0.001* | 1.99-5.60 | 2.71 | 0.001* | 1.79-4.11 | 2.47 | 0.001* | 1.31-4.66 |
| History of burn | No | ref |  |  | ref |  |  | ref |  |  |
|  | Yes | 0.78 | 0.381 | 0.43-1.44 | 1.32 | 0.062 | 0.98-1.76 | 0.91 | 0.692 | 0.55-1.52 |
| History of malnutrition | No | ref |  |  | ref |  |  | ref |  |  |
|  | Yes | 0.75 | 0.345 | 0.39-1.44 | 0.84 | 0.593 | 0.42-1.70 | 1.77 | 0.066 | 0.95-3.28 |
| History of nutritional supplementation | No | ref |  |  | ref |  |  | ref |  |  |
|  | Yes | 0.48 | 0.250 | 0.12-1.89 | 0.95 | 0.920 | 0.28-3.20 | 1.33 | 0.410 | 0.62-2.86 |

**Table S3. Adjusted odds ratios for socio-demographic and other risk factors potentially associated with skin infections in global multivariable logistic regression models including all variables**. All values were corrected for cluster sampling design.

ref = reference category used; NA = regression analysis not possible due to too few participants; *significant at p<0.05; **significant at p<0.001; †Likelihood ratio test for inclusion of the variable in the model

|  |  | **Scabies** | | | | **Pyoderma** | | | | **Fungal** | | | |
| --- | --- | --- | --- | --- | --- | --- | --- | --- | --- | --- | --- | --- | --- |
|  |  | **OR** | **p value** | **95% CIs** | **LR test†** | **OR** | **p value** | **95% CIs** | **LR test†** | **OR** | **p value** | **95% CIs** | **LR test†** |
| Sex | Male | ref |  |  |  | ref |  |  |  | ref |  |  |  |
|  | Female | 0.69 | 0.001 | 0.59-0.82 | 0.0174 | 1.21 | 0.188 | 0.89-1.64 | 0.2001 | 0.45 | 0.001 | 0.30-0.65 | <0.0001 |
| Age category | <1 year | ref |  |  |  | ref |  |  |  | ref |  |  |  |
|  | 1-2 years | 0.92 | 0.659 | 0.60-1.40 |  | 1.81 | 0.197 | 0.68-4.81 |  | 0.43 | 0.001 | 0.28-0.65 |  |
|  | 2-3 years | 0.81 | 0.545 | 0.37-1.77 |  | 2.08 | 0.103 | 0.83-5.22 |  | 0.45 | 0.006 | 0.26-0.74 |  |
|  | 3-4 years | 0.90 | 0.766 | 0.40-2.02 | 0.9314 | 2.27 | 0.148 | 0.70-7.36 | 0.1922 | 0.62 | 0.225 | 0.27-1.43 | 0.0681 |
| Tribe | Mandinka | ref |  |  |  | ref |  |  |  | ref |  |  |  |
|  | Wolof | 1.33 | 0.103 | 0.93-1.90 |  | 1.08 | 0.715 | 0.67-1.74 |  | 0.74 | 0.367 | 0.36-1.52 |  |
|  | Fula | 1.39 | 0.168 | 0.84-2.30 |  | 1.20 | 0.580 | 0.57-2.52 |  | 0.73 | 0.071 | 0.51-1.03 |  |
|  | Jola | 1.11 | 0.79 | 0.57-2.18 |  | 1.14 | 0.741 | 0.46-2.83 |  | 0.58 | 0.341 | 0.17-2.01 |  |
|  | Serehule | 1.93 | 0.035 | 1.06-3.50 |  | 1.90 | 0.046 | 1.01-3.55 |  | 0.58 | 0.229 | 0.22-1.52 |  |
|  | Other | 0.47 | 0.036 | 0.23-0.94 | 0.1089 | 0.83 | 0.663 | 0.31-2.19 | 0.3841 | 0.30 | 0.068 | 0.08-1.12 | 0.2776 |
| Mean household size | | 1.02 | 0.202 | 0.99-1.05 | 0.2617 | 1.04 | 0.013 | 1.01-1.07 | 0.0130 | 1.01 | 0.608 | 0.98-1.03 | 0.7919 |
| Mother’s education† | None | ref |  |  |  | ref |  |  |  | ref |  |  |  |
|  | Arabic school only | 0.58 | 0.029 | 0.37-0.93 |  | 1.05 | 0.913 | 0.42-2.59 |  | 0.41 | 0.020 | 0.20-0.84 |  |
|  | Primary only | 0.89 | 0.685 | 0.47-1.68 |  | 0.85 | 0.567 | 0.46-1.58 |  | 0.85 | 0.468 | 0.53-1.38 |  |
|  | Secondary only | 1.08 | 0.693 | 0.70-1.68 |  | 0.84 | 0.375 | 0.54-1.30 |  | 0.65 | 0.094 | 0.38-1.10 |  |
|  | Higher education | 1.51 | 0.321 | 0.61-3.73 | 0.3484 | 1.40 | 0.325 | 0.67-2.95 | 0.6973 | 0.57 | 0.457 | 0.11-3.02 | 0.1443 |
| Currently breastfeeding | No | ref |  |  |  | ref |  |  |  | ref |  |  |  |
|  | Yes | 1.69 | 0.024 | 1.09-2.61 | 0.0855 | 0.68 | 0.341 | 0.28-1.64 | 0.2085 | 0.50 | 0.063 | 0.24-1.05 | 0.1243 |
| Low birth weight (<2.5kg) | No | ref |  |  |  | ref |  |  |  | ref |  |  |  |
|  | Yes | 1.43 | 0.046 | 1.01-2.03 |  | 1.22 | 0.484 | 0.65-2.31 |  | 1.86 | 0.002 | 1.36-2.55 |  |
|  | Unknown | 1.22 | 0.227 | 0.86-1.73 | 0.3203 | 1.18 | 0.113 | 0.95-1.47 | 0.5205 | 1.04 | 0.765 | 0.77-1.41 | 0.2412 |
| Water source | Tap | ref |  |  |  | ref |  |  |  | ref |  |  |  |
|  | Borehole | 1.24 | 0.326 | 0.77-1.98 |  | 1.00 | 0.994 | 0.30-3.33 |  | 1.14 | 0.733 | 0.48-2.71 |  |
|  | Well | 1.21 | 0.462 | 0.67-2.12 | 0.6726 | 1.29 | 0.551 | 0.50-3.33 | 0.5373 | 1.08 | 0.754 | 0.63-1.85 | 0.9491 |
| Water distance | Inside compound | ref |  |  |  | ref |  |  |  | ref |  |  |  |
|  | <5 mins away | 0.42 | 0.075 | 0.16-1.12 |  | 0.88 | 0.706 | 0.40-1.91 |  | 1.21 | 0.565 | 0.58-2.56 |  |
|  | 5-10 mins away | 0.91 | 0.724 | 0.50-1.64 |  | 1.10 | 0.710 | 0.63-1.92 |  | 0.88 | 0.317 | 0.66-1.17 |  |
|  | >10 mins away | 1.22 | 0.687 | 0.41-3.66 | 0.2097 | 0.60 | 0.239 | 0.23-1.52 | 0.5685 | 0.45 | 0.091 | 0.17-1.17 | 0.4767 |
| Full body wash | Every day | ref |  |  |  | ref |  |  |  | ref |  |  |  |
|  | Not every day | 0.66 | 0.443 | 0.20-2.19 | 0.6878 | 2.08 | 0.231 | 0.57-7.65 | 0.5488 | NA | NA | NA | NA |
| Clean clothes | Every day | ref |  |  |  | ref |  |  |  | ref |  |  |  |
|  | Not every day | 0.44 | 0.320 | 0.07-2.64 | 0.4032 | NA | NA | NA | NA | 13.14 | 0.003 | 3.31-52.15 | 0.0001 |
| Clothes ironed | Never | ref |  |  |  | ref |  |  |  | ref |  |  |  |
|  | Sometimes | 0.93 | 0.554 | 0.73-1.20 |  | 0.96 | 0.755 | 0.72-1.29 |  | 1.28 | 0.148 | 0.90-1.84 |  |
|  | Always | 0.23 | 0.030 | 0.06-0.83 | 0.0607 | NA | NA | NA | NA | 2.10 | 0.285 | 0.47-9.39 | 0.3031 |
| Handwashing area in compound | No | ref |  |  |  | ref |  |  |  | ref |  |  |  |
|  | Yes | 0.77 | 0.079 | 0.57-1.04 | 0.1167 | 0.89 | 0.524 | 0.58-1.35 | 0.4503 | 0.83 | 0.277 | 0.58-1.19 | 0.3878 |
| Open fire in compound | No | ref |  |  |  | ref |  |  |  | ref |  |  |  |
|  | Yes | 1.59 | 0.015 | 1.12-2.26 | 0.0041 | 1.26 | 0.051 | 1.00-1.58 | 0.1434 | 1.32 | 0.386 | 0.66-2.65 | 0.1720 |
| Previous skin infection | None | ref |  |  |  | ref |  |  |  | ref |  |  |  |
|  | One | 3.14 | <0.001 | 2.12-4.65 |  | 1.95 | 0.002 | 1.40-2.72 |  | 2.41 | 0.002 | 1.57-3.71 |  |
|  | More than one | 4.86 | <0.001 | 2.83-8.33 | <0.0001 | 2.37 | 0.003 | 1.47-3.82 | <0.0001 | 2.25 | 0.040 | 1.05-4.82 | 0.0001 |
| History of burn | No | ref |  |  |  | ref |  |  |  | ref |  |  |  |
|  | Yes | 0.68 | 0.189 | 0.37-1.26 | 0.1878 | 1.01 | 0.959 | 0.63-1.63 | 0.9646 | 0.65 | 0.108 | 0.38-1.13 | 0.2182 |
| History of malnutrition | No | ref |  |  |  | ref |  |  |  | ref |  |  |  |
|  | Yes | 0.93 | 0.729 | 0.58-1.50 | 0.8194 | 0.71 | 0.339 | 0.33-1.54 | 0.2452 | 1.62 | 0.287 | 0.61-4.32 | 0.1592 |
| History of nutritional supplementation | No | ref |  |  |  | ref |  |  |  | ref |  |  |  |
|  | Yes | 0.33 | 0.243 | 0.04-2.50 | 0.1389 | 0.73 | 0.640 | 0.16-3.24 | 0.6038 | 0.59 | 0.404 | 0.15-2.33 | 0.4679 |

**Table S4. Forwards stepwise logistic regression steps.** Factor levels or variables were added to the simple model in a stepwise fashion including the next most significant, until the likelihood ratio test cutoff of >0.2 was reached. Sex and age category were included in the simple model and all subsequent steps.

*Akaike information criterion; †Likelihood ratio test for model compared to previous step.

|  | **Scabies** | | | **Pyoderma** | | | **Fungal** | | |
| --- | --- | --- | --- | --- | --- | --- | --- | --- | --- |
| **Model** | **Pseudo-R^2^** | **AIC*** | **LR test†** | **Pseudo-R^2^** | **AIC*** | **LR test†** | **Pseudo-R^2^** | **AIC*** | **LR test†** |
| Simple model | 0.0075 | 1262.503 | - | 0.0292 | 1303.954 | - | 0.0352 | 896.4075 | - |
| Step 1 | 0.0356 | 1225.17 | <0.0001 | 0.0396 | 1291.286 | 0.0001 | 0.0528 | 882.0818 | 0.0001 |
| Step 2 | 0.0538 | 1204.214 | <0.0001 | 0.0460 | 1284.711 | 0.0050 | 0.0685 | 869.4716 | 0.0001 |
| Step 3 | 0.0595 | 1199.045 | 0.0076 | 0.0508 | 1280.004 | 0.0171 | 0.0726 | 867.6772 | 0.0476 |
| Step 4 | 0.0638 | 1195.703 | 0.0203 | 0.0534 | 1278.547 | 0.0229 | 0.0753 | 867.1711 | 0.0947 |
| Step 5 | 0.0667 | 1193.965 | 0.0530 | 0.0580 | 1274.377 | 0.0434 | 0.0779 | 866.7833 | 0.1254 |
| Step 6 | 0.0692 | 1192.594 | 0.0926 | 0.0601 | 1273.316 | 0.1241 | 0.0807 | 865.9998 | 0.1487 |
| Step 7 | 0.0715 | 1191.592 | 0.0807 | 0.0624 | 1268.976 | 0.1739 | 0.0841 | 864.9258 | 0.0837 |
| Step 8 | 0.0738 | 1190.768 | 0.0926 |  |  |  | 0.0869 | 864.1161 | 0.1434 |
| Step 9 | 0.0761 | 1189.847 | 0.0887 |  |  |  |  |  |  |
| Step 10 | 0.0780 | 1189.15 | 0.1093 |  |  |  |  |  |  |
| Step 11 | 0.0798 | 1188.854 | 0.1282 |  |  |  |  |  |  |

**Table S5. Backwards elimination stepwise logistic regression steps.** Factor levels or variables were removed from the global model in a stepwise fashion removing the next least significant, until the likelihood ratio test cutoff of <0.2 was reached. Sex and age category were included in all models and steps.

*Akaike information criterion; †Likelihood ratio test for model compared to previous step.

|  | **Scabies** | | | **Pyoderma** | | | **Fungal** | | |
| --- | --- | --- | --- | --- | --- | --- | --- | --- | --- |
| **Model** | **Pseudo-R^2^** | **AIC*** | **LR test†** | **Pseudo-R^2^** | **AIC*** | **LR test†** | **Pseudo-R^2^** | **AIC*** | **LR test†** |
| Global model | 0.0866 | 1209.772 | - | 0.0694 | 1293.344 | - | 0.1014 | 880.4532 | - |
| Step 1 | 0.0865 | 1207.813 | 0.8405 | 0.0694 | 1291.344 | 0.9546 | 0.1014 | 878.4744 | 0.8886 |
| Step 2 | 0.0865 | 1205.862 | 0.8248 | 0.0693 | 1289.345 | 0.9103 | 0.1014 | 876.4775 | 0.8626 |
| Step 3 | 0.0864 | 1204.305 | 0.7822 | 0.0693 | 1287.347 | 0.9081 | 0.1006 | 879.3588 | 0.8310 |
| Step 4 | 0.0863 | 1202.416 | 0.7388 | 0.0688 | 1286.043 | 0.8096 | 0.1005 | 877.4963 | 0.8022 |
| Step 5 | 0.0862 | 1200.566 | 0.7022 | 0.0688 | 1284.122 | 0.7288 | 0.1000 | 875.9853 | 0.5190 |
| Step 6 | 0.086 | 1198.757 | 0.6615 | 0.0686 | 1282.277 | 0.6985 | 0.0992 | 874.7197 | 0.4525 |
| Step 7 | 0.0855 | 1197.373 | 0.4236 | 0.0684 | 1280.591 | 0.6089 | 0.0986 | 873.2047 | 0.4674 |
| Step 8 | 0.0849 | 1196.424 | 0.3718 | 0.0683 | 1278.799 | 0.5958 | 0.0979 | 871.8654 | 0.4315 |
| Step 9 | 0.0843 | 1195.216 | 0.373 | 0.0680 | 1277.121 | 0.5951 | 0.0970 | 870.6887 | 0.3588 |
| Step 10 | 0.0836 | 1194.06 | 0.3654 | 0.0677 | 1275.505 | 0.5213 | 0.0962 | 869.3877 | 0.3587 |
| Step 11 | 0.083 | 1192.889 | 0.3647 | 0.0674 | 1273.987 | 0.4817 | 0.0959 | 867.6681 | 0.3328 |
| Step 12 | 0.0824 | 1191.689 | 0.3799 | 0.0671 | 1272.358 | 0.5104 | 0.0949 | 866.5976 | 0.3485 |
| Step 13 | 0.0817 | 1190.48 | 0.3729 | 0.0664 | 1271.196 | 0.4286 | 0.0937 | 865.7386 | 0.3021 |
| Step 14 | 0.0808 | 1189.585 | 0.2769 | 0.0656 | 1270.278 | 0.3858 | 0.0922 | 865.1465 | 0.2384 |
| Step 15 | 0.0798 | 1188.854 | 0.2635 | 0.0649 | 1269.218 | 0.3603 | 0.0905 | 864.8105 | 0.2004 |
| Step 16 |  |  |  | 0.0639 | 1268.596 | 0.2420 | 0.0887 | 864.5341 | 0.2078 |
| Step 17 |  |  |  | 0.0624 | 1268.976 | 0.2077 | 0.0869 | 864.1161 | 0.2117 |

**Figure S3.** ***Staphylococcus aureus* isolate antibiotic sensitivities**

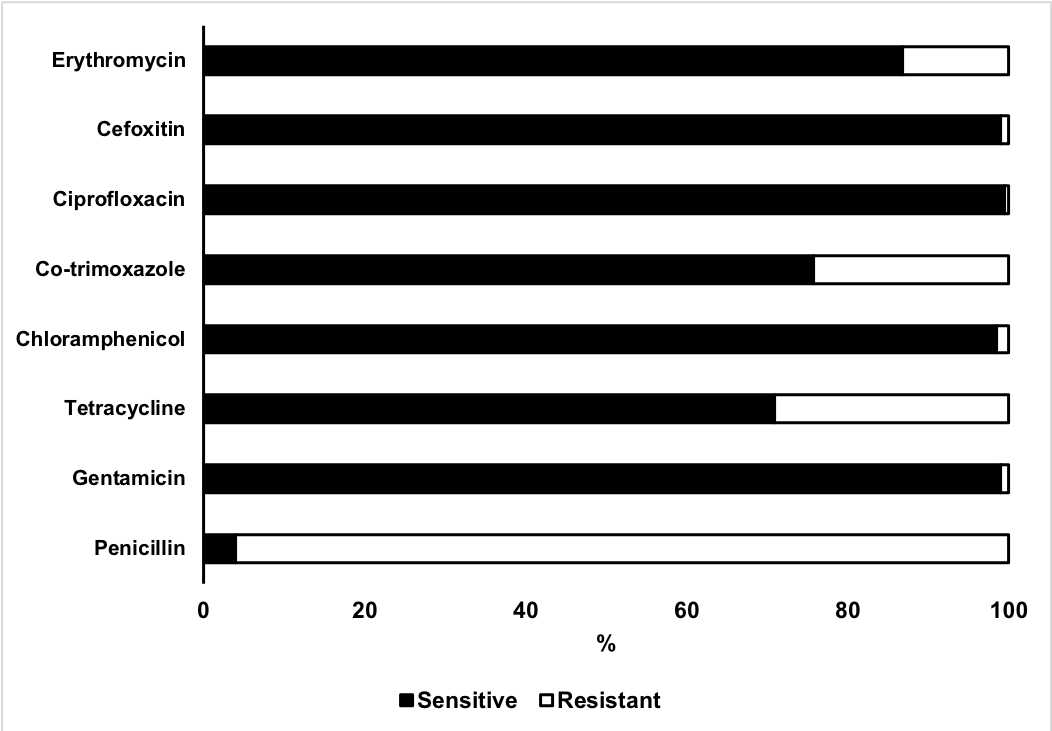

**Figure S4. Group A streptococcus isolate antibiotic sensitivities**

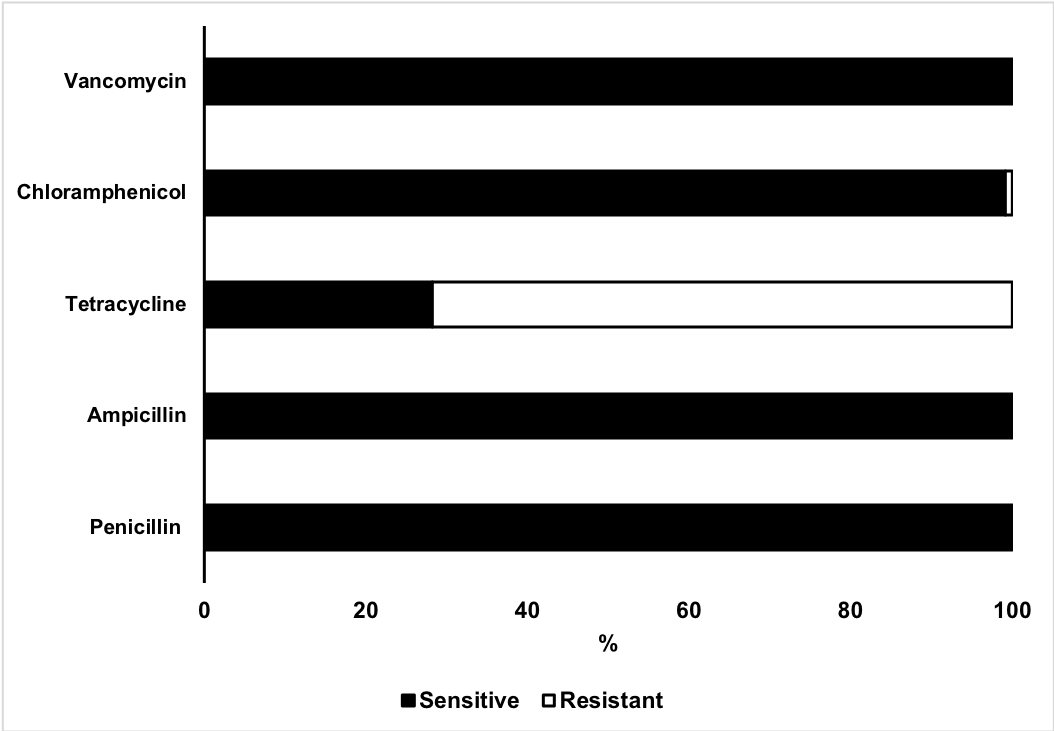

**Table S6.** **Prevalence rate ratios for presence of skin infections by whether examined before or after the start of the rainy season using Poisson regression**.

ref = reference category used; PR = prevalence ratio; †Adjusted for sex, age group, tribe, household size and mother’s education; corrected for cluster sampling design; *significant at p<0.05; **significant at p<0.001

|  |  | **Scabies** | | | **Pyoderma** | | | **Fungal** | | |
| --- | --- | --- | --- | --- | --- | --- | --- | --- | --- | --- |
|  |  | **PR** | **p value** | **95% CI** | **PR** | **p value** | **95% CI** | **PR** | **p value** | **95% CI** |
| Full sample (n=1441) | |  |  |  |  |  |  |  |  |  |
| Start of rains | Before (n=575) | ref |  |  | ref |  |  |  |  |  |
|  | After (n=866) | 1.08† | 0.702 | 0.70-1.67 | 2.42† | 0.006* | 1.38-4.23 | 0.44† | <0.001** | 0.32-0.60 |
| First / last cluster only (n=207) | |  |  |  |  |  |  |  |  |  |
| Start of rains | Before (n=101) | ref |  |  |  |  |  |  |  |  |
|  | After (n=106) | 1.59 | 0.370 | 0.58-4.37 | 2.74 | 0.014* | 1.23-6.12 | 0.60 | 0.363 | 0.19-1.82 |
|  |  | ***S. aureus* positive pyoderma** | | | **GAS positive pyoderma** | | |  |  |  |
|  |  | **PR** | **p value** | **95% CI** | **PR** | **p value** | **95% CI** |  |  |  |
| All pyoderma (n=250) | |  |  |  |  |  |  |  |  |  |
| Start of rains | Before (n=50) | ref |  |  | ref |  |  |  |  |  |
|  | After (n=200) | 1.07† | 0.571 | 0.81-1.42 | 0.99† | 0.913 | 0.75-1.30 |  |  |  |

**Table S7. Sensitivity, specificity and kappa statistic for each skin diagnosis made by trained nurses using the adapted skin condition diagnostic algorithm as compared with the diagnosis of a physician.** Study undertaken in a subset of 124 participants. ^As determined by the physician; *Kappa ranges from -1 to 1, generally values greater than >0.75 - excellent agreement, 0.4 to 0.75 - fair to good agreement, 0.4 indicate moderate or poor agreement; †Diagnosis of non-infected scabies and pyoderma, or infected scabies

|  | **Prevalence (%)^** | **Sensitivity (%)** | **Specificity (%)** | **Kappa statistic*** |
| --- | --- | --- | --- | --- |
| No skin problem | 44.4 | 98.2 | 94.2 | 0.92 |
| Non-infected scabies | 19.4 | 83.3 | 97.0 | 0.82 |
| Pyoderma | 28.2 | 97.1 | 96.6 | 0.92 |
| Infected scabies | 6.5 | 62.5 | 98.3 | 0.65 |
| Scabies and pyoderma (any combination)† | 12.9 | 81.3 | 97.2 | 0.78 |
| Fungal infection | 4.8 | 66.7 | 95.8 | 0.50 |
